# Analysis of Arabidopsis *venosa4-0* supports the role of VENOSA4 in dNTP homeostasis

**DOI:** 10.1101/2023.03.30.534755

**Authors:** Raquel Sarmiento-Mañús, Rebeca González-Bayón, Sara Fontcuberta-Cervera, Matthew A. Hannah, Francisco Javier Álvarez-Martínez, Enrique Barrajón-Catalán, Vicente Micol, Víctor Quesada, María Rosa Ponce, José Luis Micol

**Affiliations:** Instituto de Bioingeniería, Universidad Miguel Hernández, Elche, Spain; Max-Planck-Institut für Molekulare Pflanzenphysiologie, Potsdam, Germany; Instituto de Investigación, Desarrollo e Innovación en Biotecnología Sanitaria de Elche, Universidad Miguel Hernández, Elche, Spain

**Keywords:** dNTP homeostasis, Arabidopsis, *VENOSA4* gene, *ven4-0* mutant, SAMHD1 ortholog

## Abstract

An imbalance in the deoxyribonucleoside triphosphate (dNTP) pool caused by an increase or decrease in the levels of any of the four dNTPs leads to increased DNA mutations, overloading DNA repair mechanisms. The human protein SAMHD1 (Sterile alpha motif and histidine-aspartate domain containing protein 1) functions as a dNTPase to maintain the balance of the dNTP pool, as well as in DNA repair. In eukaryotes, the limiting step in *de novo* dNTP synthesis is catalyzed by RIBONUCLEOTIDE REDUCTASE (RNR), which consists of two R1 and two R2 subunits. In Arabidopsis, RNR1 is encoded by *CRINKLED LEAVES 8* (*CLS8*) and RNR2 by three paralogous genes, including *TSO2* (*TSO MEANING ’UGLY’ IN CHINESE 2*). In plants, the *de novo* biosynthesis of purines occurs within the chloroplast, and DOV1 (DIFFERENTIAL DEVELOPMENT OF VASCULAR ASSOCIATED CELLS 1) catalyzes the first step of this pathway. Here, to explore the role of *VENOSA4* (*VEN4*), the most likely Arabidopsis ortholog of human *SAMHD1*, we studied the *ven4-0* mutant. The mutant leaf phenotype caused by the *ven4-0* point mutation was stronger than those of T-DNA insertional *ven4* mutations. Structural predictions suggested that the E249L amino acid substitution in the mutated VEN4-0 protein rigidifies its 3D structure compared to wild-type VEN4. The morphological phenotypes of the *ven4*, *cls8*, and *dov1* single mutants were similar, and those of the *ven4 tso2* and *ven4 dov1* double mutants were synergistic. The *ven4-0* mutant had reduced levels of four amino acids related to dNTP biosynthesis, including glutamine and glycine, which are precursors in the *de novo* purine biosynthesis pathway. Finally, despite its annotation in some databases, At5g40290, a paralog of *VEN4*, is likely a pseudogene. These observations support the previously proposed role of VEN4 in dNTP metabolism. Our results reveal a high degree of cross-kingdom functional conservation between VEN4 and SAMHD1 in dNTP homeostasis.

## INTRODUCTION

Deoxyribonucleoside triphosphates (dATP, dCTP, dGTP, and dTTP; collectively referred to as dNTPs hereafter) are present in the cells of all living beings, functioning as DNA precursors. Indeed, a continuous input of dNTPs is required for replication, repair, and recombination of the nuclear, mitochondrial, and chloroplastic (in photosynthetic organisms) genomes. Two dNTP synthesis pathways have been described in all eukaryotes and most prokaryotes: the *de novo* and salvage (recycling) pathways (Kilstrup et al., 2005; Guarino et al., 2014; Witte and Herde, 2020). In eukaryotes, the limiting step of the *de novo* pathway is catalyzed by RIBONUCLEOTIDE REDUCTASE (RNR), which consists of two R1 (also called α) major subunits and two R2 (β) minor subunits (Jordan and Reichard, 1998). Transcription of the genes encoding RNR subunits is activated in the S phase of the cell cycle, as well as in response to DNA damage (Guarino et al., 2014).

In *Arabidopsis thaliana* (hereafter, Arabidopsis), the RNR R1 subunit is encoded by a single gene: *RNR1*, also known as *CRINKLED LEAVES 8* (*CLS8*; Garton et al., 2007) and *DEFECTIVE IN POLLEN ORGANELLE DNA DEGRADATION 2* (*DPD2*; Tang et al., 2012). In the *cls8-1* mutant, the first two leaves are yellowish, and the remaining leaves are wrinkled with irregular, whitish margins; the flowers are asymmetrical, with wrinkled petals (Garton et al., 2007). Three paralogous genes encode the RNR R2 subunit in Arabidopsis: *TSO MEANING ’UGLY’ IN CHINESE 2* (*TSO2*), *RNR2A*, and *RNR2B*; TSO2 is the major contributor to RNR function. In the *tso2-1* mutant, leaves from the fifth and subsequent nodes display whitish areas and irregular margins. Some *tso2-1* plants also show fasciated stems, homeotic transformations of floral organs, and reduced fertility. Although the *rnr2a-1* and *rnr2b-1* single mutants and the *rnr2a-1 rnr2b-1* double mutant appear phenotypically wild type, *tso2-1 rnr2a-1* and *tso2-1 rnr2b-1* are lethal (Wang and Liu, 2006), providing evidence for the functional redundancy of TSO2 with RNR2A and RNR2B. As expected, dNTP levels are substantially reduced in the *cls8-1* and *tso2-1* mutants due to their loss of RNR activity (Wang and Liu, 2006; Garton et al., 2007).

Human Sterile alpha motif and histidine-aspartate domain containing protein 1 (SAMHD1) is a nuclear protein with dGTP-dependent triphosphohydrolase activity, which, in contrast to RNR, degrades dNTPs into deoxyribonucleosides and inorganic triphosphate. SAMHD1 harbors a histidine (H) and aspartic acid (D)-rich (HD) domain, which is required for its dNTPase activity (Aravind and Koonin, 1998), and a Sterile alpha motif (SAM), whose function is unclear but might help stabilize SAMHD1 during antiviral activity (Stillman, 2013; Shigematsu et al., 2014; Mauney and Hollis, 2018). Mutations in *SAMHD1* have been associated with Aicardi-Goutières syndrome, a congenital neurodegenerative autoimmune disorder with early childhood onset and symptoms similar to those of a congenital viral infection (Goldstone et al., 2011; Powell et al., 2011; Kretschmer et al., 2015). SAMHD1 appears to also be involved in defense against viruses, as it degrades dNTPs following infection by human immunodeficiency virus (HIV), thus hindering the reverse transcription of the viral genome (Kretschmer et al., 2015). Mutations in *SAMHD1* have also been detected in some types of cancer (Li et al., 2017; Coggins et al., 2020).

Putative orthologs of SAMHD1 have been studied in Arabidopsis (VENOSA4 [VEN4]) and *Oryza sativa* (rice; STRIPE3 [ST3]), and are related to chloroplast and leaf development, stress responses, and dNTP metabolism (Yoshida et al., 2018; Xu et al., 2020; Wang et al., 2022). VEN4 hydrolyzes dGTP to 2′-deoxyguanosine (2′-dG) *in vitro* and positively regulates plant immunity (Lu et al., 2022). Here, to further explore the roles of *VEN4* and its genetic interactions and possible involvement in dNTP metabolism, we studied three *ven4* allelic mutants. Our analysis of these mutants, particularly the original *ven4-0* point mutation, revealed a high degree of cross-kingdom functional conservation between Arabidopsis *VEN4* and its likely human ortholog *SAMHD1*.

## MATERIALS AND METHODS

### Plant materials, growth conditions, and genotyping

The *Arabidopsis thaliana* (L.) Heynh. wild-type accessions Col-0 and L*er* and the *ven4-2* (SALK_077401), *ven4-3* (SALK_131986), *rnr2a-2* (SALK_150365), and SALK_121024 mutants in the Col-0 genetic background were obtained from the Nottingham Arabidopsis Stock Centre (NASC). The *ven4-0* mutant in the L*er* background was isolated in the laboratory of J.L. Micol and was previously described as *ven4* (Berná et al., 1999; Robles and Micol, 2001; Bensmihen et al., 2008; Pérez-Pérez et al., 2011). Seeds of *tso2-1* (in the L*er* background) were kindly provided by Zhongchi Liu (University of Maryland, College Park, MD, USA), and *dov1* seeds (in the En-2 background) by Kevin Pyke (University of Nottingham, Sutton Bonington, Leicestershire, UK). Plants were grown under sterile conditions on half-strength Murashige and Skoog (MS; Duchefa Biochemie) medium containing 0.7% plant agar (Duchefa Biochemie) and 1% sucrose (Duchefa Biochemie) at 20°C ± 1°C, 60-70% relative humidity, and under continuous fluorescent light of ≈75 μmol/m^2^·s and crossed as previously described (Ponce et al., 1998; Berná et al., 1999). Unless otherwise stated, all plants used were homozygous for the mutations indicated. Mapping of *ven4-0* and genotyping of single and double mutants was made by PCR amplification and/or Sanger sequencing using the primers described in Supplementary Tables S1 and S2.

### Phenotypic and morphometric analyses and microscopy

Rosettes, siliques, stems, and inflorescences were photographed using a Leica MZ6 stereomicroscope equipped with a Nikon DXM1200 digital camera. Light microscopy, confocal imaging, and transmission electron microscopy were performed as previously described (Quesada et al., 2011). The NIS Elements AR 3.1 image analysis package (Nikon) was used to measure rosette area and hypocotyl length. Main stem length was measured with a ruler.

### Chlorophyll concentration, photosynthetic efficiency, and fresh and dry weight measurements

Chlorophyll concentration in μg per ml of plant extract was measured as previously described (Lichtenthaler and Wellburn, 1983), as [chlorophyll a] = 12.21·A_663_ − 2.81·A_646_, and [chlorophyll b] = 20.31·A_646_ − 5.03·A_663_. The chlorophyll content was then recalculated on a plant fresh-weight basis as μg of chlorophyll a or b per mg of plant fresh weight (μg/mg). Photosynthetic yield was measured in the central region of the lamina of the third-node leaf of each seedling using a DUAL-PAM-100 portable chlorophyll fluorometer (WALZ, Effeltrich, Germany) immediately after 30 min of dark adaption. The fresh weights of the seedlings were measured immediately after collection, and dry weights were measured after drying overnight in an oven at 55°C.

### Construction of transgenic lines

For transgenic complementation of the *ven4-0*, *ven4-2*, and *ven4-3* mutations, a 7-kb region extending from the nucleotide 3,416 upstream of the translation start codon to the last nucleotide of the 3′-UTR of *VEN4* was PCR amplified using the primer pair VEN4pro:VEN4_F/R (Supplementary Table S2). The PCR amplification products were cloned into the pGreenII0179 vector (Hellens et al., 2000) after restriction with *Not*I and *Sal*I and ligation with T4 ligase (Fermentas).

To obtain the *VEN4_pro_:GUS* and *35S_pro_:VEN4:GFP* constructs, the 3,416-bp genomic region upstream of the translation start codon of *VEN4* or the full-length coding sequence of *VEN4* (with the translation stop codon removed to obtain GFP translational fusions) was PCR amplified with primer pairs VEN4pro:GUS_F/R and 35Spro:VEN4:GFP_F/R, respectively, as described in Supplementary Table S2. The PCR amplification products were cloned into the pENTR/D-TOPO Gateway entry vector (Invitrogen) via BP reactions. The *VEN4_pro_* and *VEN4* (without its stop codon) inserts of the entry clones were subcloned into the pMDC164 and pMDC83 destination vectors, respectively, via LR reactions (Curtis and Grossniklaus, 2003).

Chemically competent *Escherichia coli* DH5α cells were transformed by the heat-shock method with the ligation products or the BP and LR reaction mixes. The integrity of the transgenes was verified by Sanger sequencing of at least two independent transformant clones. *Agrobacterium tumefaciens* LBA4404 cells were transformed by electroporation with the verified constructs. The transgenes were transferred into Col-0 (*VEN4_pro_:GUS* and *35S_pro_:VEN4:GFP*) or *ven4-0*, *ven4-2*, and *ven4-3* plants (*VEN4_pro_:VEN4*) by the floral dip method (Clough and Bent, 1998). The *ven4-0* and *ven4-2* mutants were crossed to Col-0 *35S_pro_:VEN4:GFP* plants, and 8 F_2_ Hyg^R^ plants were genotyped to identify *ven4* homozygotes carrying at least a single copy of the transgene.

### RNA isolation, RT-PCR, and RT-qPCR

For gene expression analysis, RNA was isolated from the aerial parts of plants collected 15 or 21 days after stratification (das) using TRIzol (Invitrogen). cDNA synthesis and PCR amplifications were carried out as previously described (Wilson-Sánchez et al., 2018). qPCR amplification was carried out in a Step-One Real-Time PCR System (Applied Biosystems), with three technical replicates per biological replicate (each consisting of three rosettes). The primers used are described in Supplementary Table S2. The housekeeping gene *ACTIN2* (*ACT2*) was used as an internal control for relative quantification, as previously described (Wilson-Sánchez et al., 2018). The C_T_ values were normalized using the 2^-ΔΔC^_T_ method (Livak and Schmittgen, 2001).

### Metabolite profiling

Third- and fourth-node leaves were collected 21 das from at least six biological replicates of *ven4-0* and L*er* plants. Metabolite profiling was performed by GC-MS as previously described (Lisec et al., 2006). Targeted metabolite identification was performed using the TargetSearch Bioconductor package (Cuadros-Inostroza et al., 2009) with a library based on approximately 900 reference compounds from the GMD database (Kopka et al., 2005; Schauer et al., 2005). Only compounds with a retention index (RI) deviation of <2000 and an identification based on at least five matching correlated masses were retained. Redundant metabolites were either removed or grouped to retain the most likely identification, and other potential hits were noted. Metabolites with significant differences between L*er* and *ven4-0* were identified based on a Student’s *t*-test.

### Protein structure visualization and analysis

The 3D structures of the full-length monomers of VEN4 and human SAMHD1 were downloaded from the AlphaFold Protein Structure Database (AlphaFold DB; Jumper et al., 2021; Varadi et al., 2021; https://alphafold.ebi.ac.uk; in this database, VEN4 and human SAMHD1 are identified as AF-Q9FL05-F1 and AF-Q9Y3Z3-F1, respectively) and visualized using the UCSF ChimeraX 1.2.5 software (Goddard et al., 2018; Pettersen et al., 2021; https://www.rbvi.ucsf.edu/chimerax/).

To analyze the impact of the E249L substitution on the conformational stability and dynamics of VEN4 and the equivalent E355L mutation in human SAMHD1, we used two web structure-based protein stability predictors: DynaMut (Rodrigues et al., 2018; https://biosig.lab.uq.edu.au/dynamut/) and DynaMut2 (Rodrigues et al., 2021; https://biosig.lab.uq.edu.au/dynamut2/). These predictors quantify the difference in the unfolding Gibbs free energy between wild-type and mutant proteins (ΔΔG, expressed in kcal/mol) and classify mutations as stabilizing when ΔΔG > 0 kcal/mol or destabilizing when ΔΔG < 0 kcal/mol. DynaMut also offers the ΔΔG results from three additional predictors: SDM (Worth et al., 2011), mCSM (Pires et al., 2014b), and DUET (Pires et al., 2014a), as well as the difference in the vibrational entropy energy between wild-type and mutant proteins (ΔΔS_Vib_, expressed in kcal/mol/K), as computed by the ENCoM server (Frappier et al., 2015), which classifies mutations as rigidifying if ΔΔS_Vib_ > 0 kcal/mol/K, or flexibilizing when ΔΔS_Vib_ < 0 kcal/mol/K. Finally, we used Missense3D (Ittisoponpisan et al., 2019; http://missense3d.bc.ic.ac.uk/missense3d/) to predict damaging structural effects on VEN4 and human SAMHD1 proteins upon E249L and E355L substitutions, respectively.

### Accession numbers

Sequence data from this article can be found at TAIR (http://www.arabidopsis.org) under the following accession numbers: *VEN4* (At5g40270), *VEN4* paralog (At5g40290), *RNR1* (At2g21790), *TSO2* (At3g27060), *RNR2A* (At3g23580), *RNR2B* (At5g40942), *DOV1* (At4g34740), and *ACT2* (At3g18780).

## RESULTS

### Positional cloning of the *ven4-0* mutation

A number of Arabidopsis mutants exhibit rosette leaf reticulation, with some, most, or all veins green but the interveinal tissues pale. This phenotype is usually due to a reduced number of interveinal mesophyll cells and/or alterations in chloroplast development, which result in locally reduced contents of chlorophylls and other photosynthetic pigments (reviewed in Lundquist et al., 2014). In a large-scale screening for ethyl methanesulfonate (EMS)-induced Arabidopsis mutants with abnormal leaf shape, size or pigmentation (Berná et al., 1999), we previously isolated hundreds of viable mutants, some of which were named *venosa* (*ven*) since they exhibited reticulated rosette leaves. One such mutation, named *ven4* (referred to here as *ven4-0*), is recessive and fully penetrant (Berná et al., 1999), has only mild effects on whole leaf shape (Bensmihen et al., 2008), and increases the number of stomata in leaves (Pérez-Pérez et al., 2011).

We subjected the *ven4-0* mutation, which was isolated in the L*er* background (Figure 1A, B), to iterative linkage analysis using molecular markers, finding that the At5g40270 gene was the best candidate to be *VEN4* (Figure 2). We crossed *ven4-0* (Figure 1B) to two lines harboring T-DNA insertions in At5g40270: SALK_077401 and SALK_131986 (in the Col-0 background; Figure 1C-E). Non-complementation was observed in the F_1_ plants of these crosses, confirming that At5g40270 is *VEN4*. We initially named these two lines *ven4-2* and *ven4-3*, respectively (Supplementary Figure S1). We later found that *phyB-9*, an extensively studied mutant line assumed to carry only a mutant allele of *PHYTOCHROME B* (*PHYB*), also harbors a mutant allele of *VEN4*, which we referred to as *bnen* in Yoshida et al. (2018); we also stated that *ven4-2* and *bnen* are alleles of *VEN4*. We mentioned in that paper that we already identified At5g40270 as *VEN4* and indicated that we would describe its identification and the analysis of other *ven4* alleles elsewhere.

**Figure 1.**
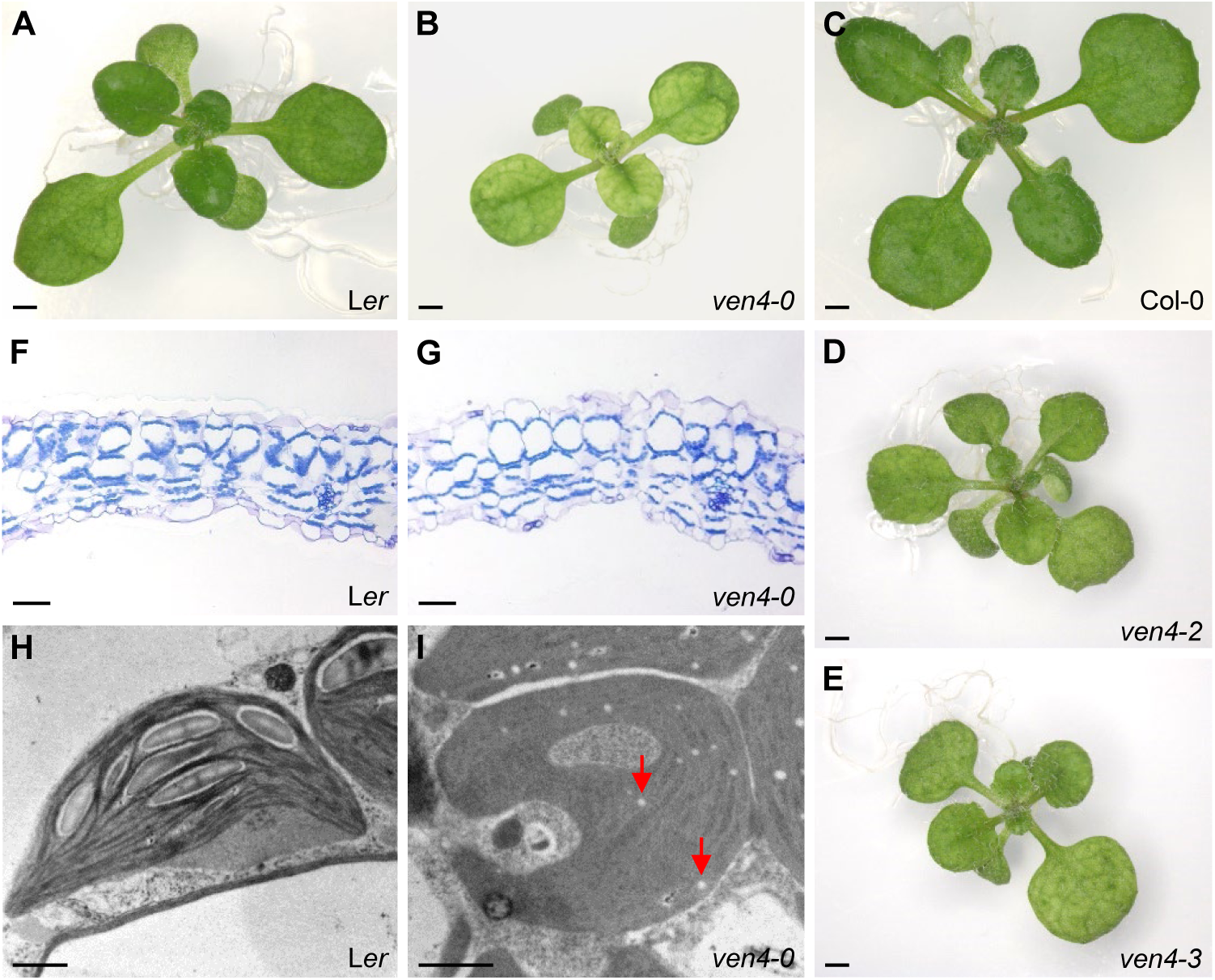
Morphological and histological phenotypes of the *ven4* alleles examined in this study. A to E, Rosettes of wild-type L*er* (A) and Col-0 (C) and the *ven4-0* (B)*, ven4-2* (D), and *ven4-3* (E) mutants. Photographs were taken 16 days after stratification (das). Scale bars: 2 mm. F and G, Transverse sections midway along the leaf margin and the primary vein of L*er* (F) and *ven4-0* (G) third-node rosette leaves. Scale bars: 40 μm. H and I, Transmission electron micrographs of L*er* (H) and *ven4-0* (I) chloroplasts. Red arrows in (I) indicate plastoglobules. Third-node leaves were collected 21 das. Scale bars: 1 µm.

**Figure 2.**
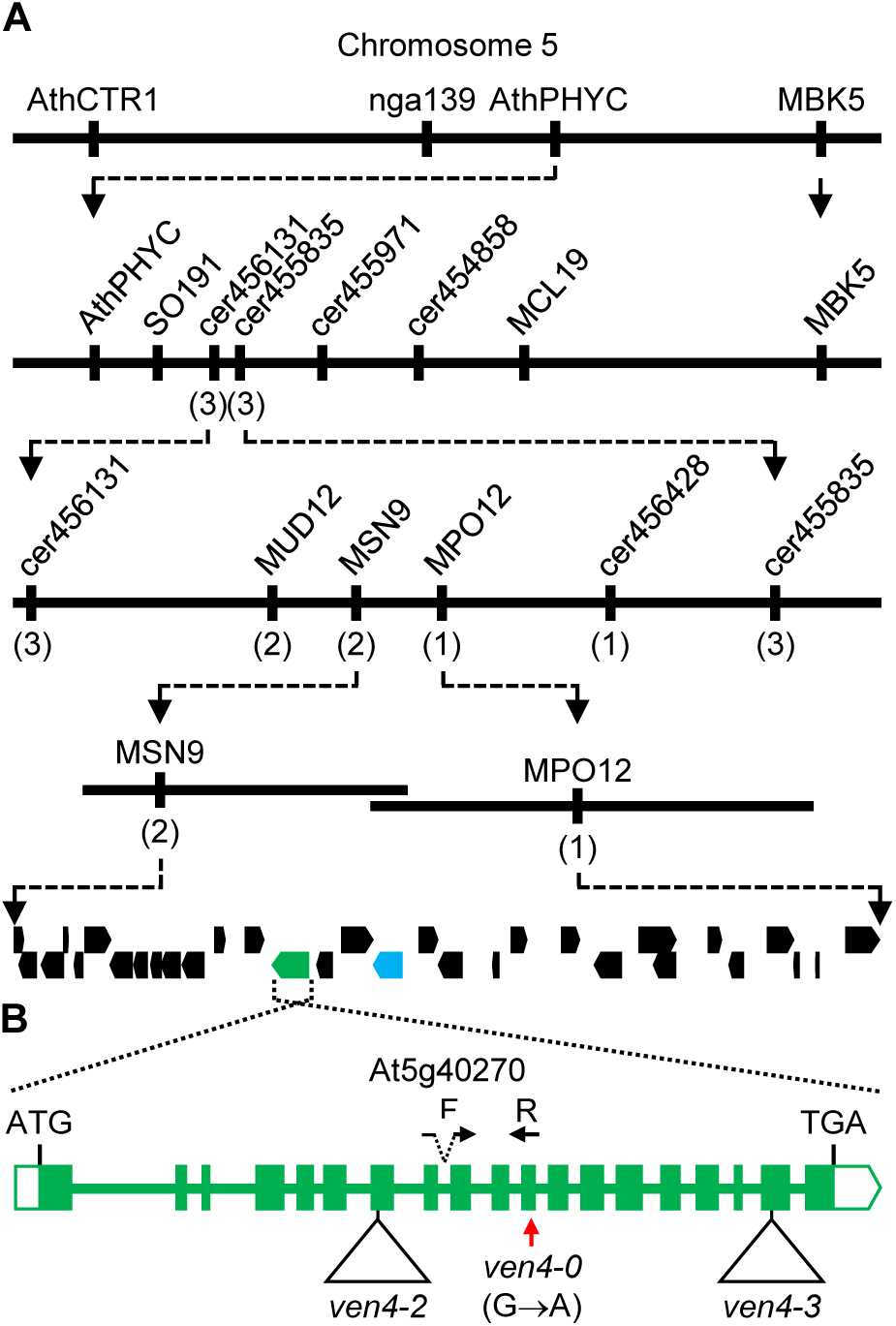
Positional cloning and structure of the *VEN4* gene. A, Map-based strategy used to identify the *VEN4* gene. Using the mapping method described in Ponce et al. (1999), Robles and Micol (2001) found the *ven4-0* mutation linked to the AthPHYC marker. The linkage analysis of 300 plants from an F_2_ mapping population derived from a cross between Col-0 and *ven4-0* allowed us to delimit a candidate interval of 100 kb (encompassing 31 genes). The molecular markers used for linkage analysis (see also Supplementary Table S1) and the number of informative recombinants identified (in parentheses) are indicated. To identify *VEN4* among the candidate genes, we searched for publicly available Salk T-DNA lines (Alonso et al., 2003; http://signal.salk.edu) carrying insertions within the interval. Two of the 33 T-DNA insertional lines that we tested displayed phenotypic traits similar to those of *ven4-0* plants, and their T-DNA insertions were shown to disrupt the seventh and eighteenth exons of At5g40270, respectively. PCR amplification and Sanger sequencing confirmed the presence and positions of the annotated insertions at nucleotide positions 1,583 (from the predicted translation start codon) in SALK_077401 and 3,409 in SALK_131986. A complementation test demonstrated allelism between *ven4-0* and these two T-DNA lines (Supplementary Figure S1). The At5g40270 gene is shown in green and its paralog At5g40290 in blue. B, Structure of the *VEN4* gene indicating the positions and nature of the *ven4* mutations examined in this study and the positions of the RT-qPCR_VEN4_F and RT-qPCR_VEN4_R primers (shown as F and R, respectively; see Supplementary Table S2), which are not drawn to scale. Exons and introns are indicated by boxes and lines, respectively; filled and open boxes represent coding sequences and untranslated regions, respectively. Triangles represent T-DNA insertions, and the vertical red arrow shows the *ven4-0* point mutation. The translation start (ATG) and stop (TGA) codons are also shown.

During the course of the current study, Xu et al. (2020) proposed that VEN4 is involved in dNTP metabolism based on (1) structural homology with human SAMHD1 and (2) the finding that treatment with dNTPs partially rescued the phenotypes of two T-DNA-insertional *ven4* mutants. The authors named the *VEN4* allele carried by the SALK_023714 line *ven4-1* (which we did not examine here) and the *VEN4* allele carried by the SALK_077401 line *ven4-2* (as also described in Yoshida et al., 2018). Hence, to avoid confusion, we introduce the names *ven4-0* to describe the EMS-induced allele that we previously referred to as *ven4* since 1999 (Berná et al., 1999; Robles and Micol, 2001; Bensmihen et al., 2008; Pérez-Pérez et al., 2011) and *ven4-3* for SALK_131986. The other alleles are referred to as previously named: *ven4-1* for SALK_023714 (Xu et al., 2020; Lu et al., 2022) and *ven4-2* for SALK_077401 (Yoshida et al., 2018; Xu et al., 2020; Lu et al., 2022).

Sanger sequencing of the *ven4-0* allele revealed a G→A transition (Figure 2B and Supplementary Table S2) that is predicted to cause an E249L missense substitution in the protein encoded by At5g40270. RT-qPCR revealed extremely low levels of *ven4-2* transcripts, including sequences downstream of its T-DNA insertion, suggesting that this allele is nearly null (its 2^-ΔΔC^_T_ is 96·10^-4^ fold that of Col-0).

### *ven4-0* exhibits stronger leaf defects compared to its T-DNA-insertional *ven4* **alleles**

The mutant morphological phenotype of *ven4-0* (Supplementary Table S3) was stronger than those of *bnen* (Yoshida et al., 2018), *ven4-1* (Xu et al., 2020) and *ven4-2* (Yoshida et al., 2018; this work). Similarly, the reductions in chlorophyll contents and photosynthetic efficiency were more pronounced in *ven4-0* plants (Supplementary Table S4). The leaf internal structure was similar in *ven4-0* and L*er* (Figure 1F, G), but the mutant had smaller chloroplasts in the palisade mesophyll cells, as revealed by transmission electron microscopy (Figure 1H, I); this phenotype was previously observed in the *bnen* mutant (Yoshida et al., 2018). In addition, *ven4-0* chloroplasts exhibited an increased number of plastoglobules, reduced number of starch granules, and poorly organized thylakoids compared to L*er* (Figure 1H, I). In agreement with the aberrant chloroplast development exhibited by *ven4* mutants, the *ven4-1* mutant exhibits markedly reduced levels of photosynthetic proteins (Xu et al., 2020). Finally, the leaf paleness of *ven4* mutants is a temperature-sensitive trait: growth at 26°C restored leaf color in *ven4-0*, *ven4-2*, and *ven4-3* to wild-type levels (Supplementary Figure S2). This result is also in agreement with the recovery of photosynthetic activity observed when *bnen* and *ven4-1* were shifted from 22°C to 32°C (Xu et al., 2020).

VEN4 is thought to play an important role in leaf chloroplast development mainly following leaf emergence (Xu et al., 2020). However, under our growth conditions, GUS activity in five independent lines carrying the *VEN4_pro_:GUS* transgene was highest in emerging leaves and decreased with leaf expansion (Figure 3A), pointing to a photosynthesis-independent function of VEN4. We observed high *VEN4* promoter activity in emerging leaves and developing flowers, suggesting that VEN4 functions in highly proliferative tissues (Figure 3C, E). We also detected GUS staining in the stem and root vasculature (Figure 3B, D).

**Figure 3.**
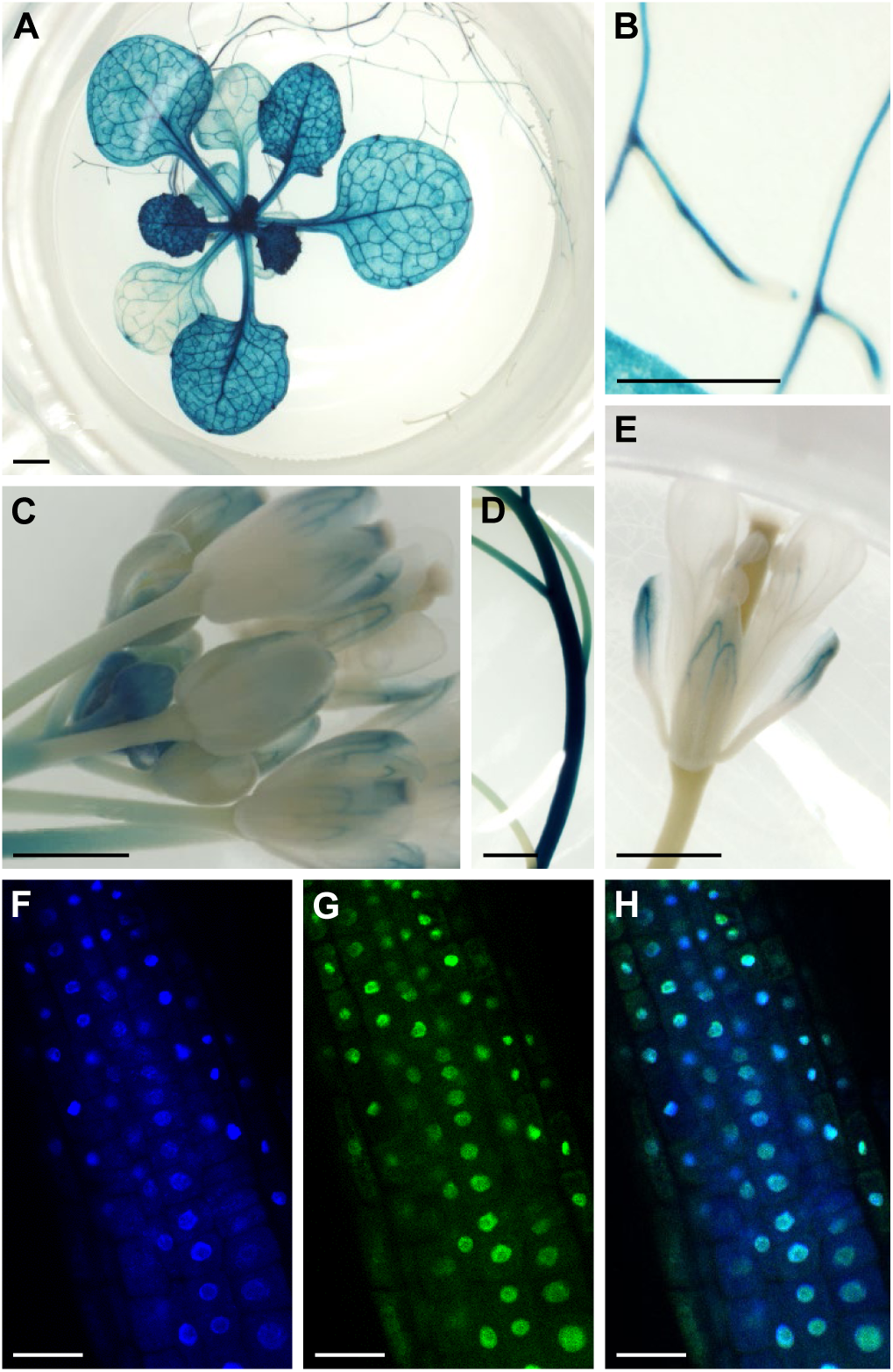
Expression pattern of *VEN4* and subcellular localization of VEN4 in wild-type Col-0. A to E, Visualization of *VEN4_pro_:GUS* transgene activity in rosette (A), root (B), inflorescence (C), stem (D), and flower tissue (E). Seedlings and plant organs were collected 21 (A and B) and 34 (C, D and E) das. Scale bars: 1 mm (A, C, and E) and 0.5 mm (B and D). F to H, Apical root cells of a *35S_pro_:VEN4:GFP* transgenic plant in the Col-0 background, showing fluorescence from 4′,6-diamidino-2-phenylindole (DAPI) (F), GFP (G), and their overlay (H). Roots were collected 6 das. Scale bars: 20 µm.

We confirmed the identity of *VEN4* by transforming *ven4-0*, *ven4-2*, and *ven4-3* with a transgene containing a 7,364-bp Col-0 genomic DNA fragment including the At5g40270 coding region (4,209 bp) and its putative promoter; this *VEN4_pro_:VEN4* transgene fully rescued the mutant phenotypes of *ven4-0*, *ven4-2*, and *ven4-3* plants (Supplementary Figure S3A-F). In addition, the expression of *35S_pro_:VEN4:GFP* fully restored the wild-type phenotypes of *ven4-0* and *ven4-2* (Supplementary Figure S3G-J). In these transgenic plants, VEN4 localized to the nucleoplasm of apical root cells in a diffuse pattern (Figure 3F-H), like human SAMHD1 (Kretschmer et al., 2015). These results confirm previous findings from Xu et al. (2020), which were obtained by transient transgene expression in Arabidopsis leaf protoplasts.

### The mutant VEN4-0 protein is predicted to be more rigid than wild-type VEN4

The morphological phenotype caused by the *ven4-0* point mutation is stronger than those of the T-DNA insertional *ven4* alleles. To explore the molecular basis of such differences in phenotypic strength, we compared the 3D structures of full-length VEN4 and human SAMHD1 proteins from AlphaFold DB (Jumper et al., 2021; Varadi et al., 2021; https://alphafold.ebi.ac.uk/).

The dNTPase activity of human SAMHD1 is regulated by the combined action of GTP and all four dNTPs, with which SAMHD1 assembles into functional tetramers. When dATP binds to allosteric site 2 of SAMHD1 (N119, D330, N358, and R372), E355 moves, enabling the establishment of a salt bridge with the R333 residue, which stabilizes the bound dATP via a stacking interaction (Ji et al., 2013; Ji et al., 2014). SAMHD1 R333 appears to be equivalent to VEN4 R231, which is also close enough to E249 (the residue affected by the *ven4-0* mutation) to form a salt bridge, as shown by our 3D-structural prediction (Figure 4B).

**Figure 4.**
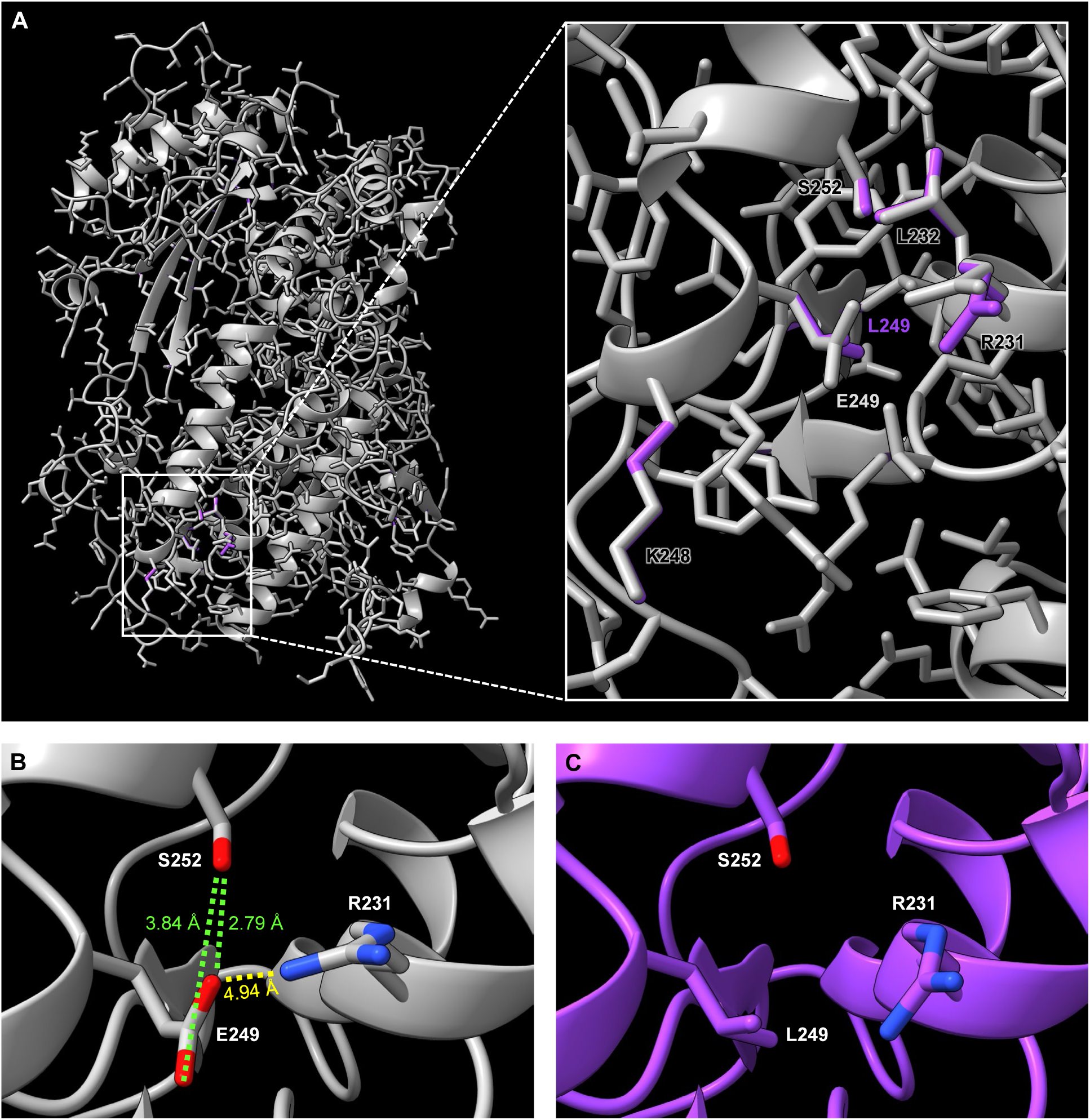
Comparison of the 3D structures of the VEN4 wild-type and E249L mutant (VEN4-0) proteins. A, Cartoon representation of the overlay of the 3D structures of VEN4 (colored in gray) and VEN4-0 (colored in purple), with a close-up view of the vicinity of the substituted E. All residues affected by the mutation, whose side chains show different angles between wild-type and mutant proteins, are labeled. B and C, Non-covalent interactions disrupted by the E249L substitution; the distances between the interacting atoms are indicated in angstroms (Å). The hydrogen bonds and salt bridges are represented by green and yellow dotted lines, respectively. The interacting oxygen and nitrogen atoms are highlighted in red and blue, respectively.

To assess the potential effects of the *ven4-0* mutation (E249L) on the structural stability and dynamics of VEN4-0, we predicted differences in the unfolding Gibbs free energy (ΔΔG) and vibrational entropy energy (ΔΔS_Vib_) between the wild-type and mutant proteins using DynaMut (Rodrigues et al., 2018) and DynaMut2 (Rodrigues et al., 2021). Both servers estimated positive ΔΔG values indicating that E249L is a stabilizing mutation (Supplementary Table S5). A negative ΔΔS_Vib_ value was also calculated by ENCoM (Frappier et al., 2015), pointing to the possible rigidification of the 3D structure of VEN4-0 compared to VEN4 (Supplementary Table S5 and Supplementary Figure S4A). We also conducted this analysis simulating an equivalent E355L mutation in human SAMHD1, which revealed that this mutation would also stabilize and rigidify the structure of SAMHD1 (Supplementary Table S5 and Supplementary Figure S4B).

We used Missense3D (Ittisoponpisan et al., 2019) to predict the damaging effects of the E249L change on VEN4-0 structure compared to VEN4 and the effects of the equivalent E355L change in SAMHD1. No structural damage was predicted by Missense3D for the E249L substitution of VEN4-0 based on changes in solvent exposure; indeed, residue 249 is exposed to solvents in a similar manner with (L249) and without (E249) the mutation. Nevertheless, when we compared the 3D structures of wild-type and mutant proteins, we observed changes in the side chain angles of some residues in the vicinity of the mutated amino acid (Figure 4A). We also detected the loss of two hydrogen bonds between E249 and S252, as well as the salt bridge between E249 and R231, whose impact was not predicted by the software mentioned above (Figure 4B, C). By contrast, Missense3D classified the equivalent E355L substitution in human SAMHD1 as structurally damaging due to the replacement of a buried negative charge, as well as the disruption of two buried hydrogen bonds and the salt bridge between the E355 and R333 residues (Supplementary Figure S5B, C). The overlay of wild-type and mutant proteins also showed differences in the side chain angles of neighboring amino acids, including R333, which is located close to E355 in the secondary but not the primary structure of SAMHD1 (Supplementary Figure S5A).

### The *ven4-0* and *dov1* mutations genetically interact and cause opposite amino acid profiles

In some Arabidopsis reticulated mutants, such as *differential development of vascular associated cells 1* (*dov1*), leaf veins appear green because the perivascular bundle sheath cells are apparently normal. The remaining mesophyll tissue shows increased numbers of air spaces and a reduced number of cells, which are malformed and, in some cases, contain morphologically aberrant chloroplasts. *DOV1* encodes glutamine phosphoribosyl pyrophosphate aminotransferase 2 (ATase2), which catalyzes the first step of purine nucleotide biosynthesis within the chloroplast (Rédei and Hirono, 1964; Li et al., 1995; Kinsman and Pyke, 1998; Mollá-Morales et al., 2011; Rosar et al., 2012). Indeed, some steps of purine and pyrimidine biosynthesis take place in chloroplasts and require several amino acids as substrates. In the *dov1* mutant, the levels of glycine, alpha-alanine, proline, asparagine, aspartate, lysine, the asparagine precursor ornithine, and inorganic phosphate are significantly increased (Hung et al., 2004).

To determine whether the functions of *VEN4* and *DOV1* are related, we generated *ven4-0 dov1* and *ven4-2 dov1* double mutants, which exhibited a synergistic phenotype: their growth was slow, and their leaves were small, irregularly shaped, and accumulated anthocyanins (Figure 5A, B, D, G, H). We then performed a metabolomic analysis of *ven4-0* and L*er*. The amino acid profile of *ven4-0* was opposite to that published for *dov1* in terms of the levels of glycine, proline, and asparagine. We also detected reduced levels of other amino acids: glutamine, phenylalanine, tyrosine, valine, beta-alanine, methionine, cysteine, and arginine (Figure 6 and Supplementary Table S6). Four of these amino acids are related to different steps of dNTP biosynthesis: glutamine is a precursor of both purines and pyrimidines; glycine is a specific precursor of purines; arginine biosynthesis shares intermediates with pyrimidine *de novo* biosynthesis; and beta-alanine is a product of uridine catabolism (Witte and Herde, 2020).

**Figure 5.**
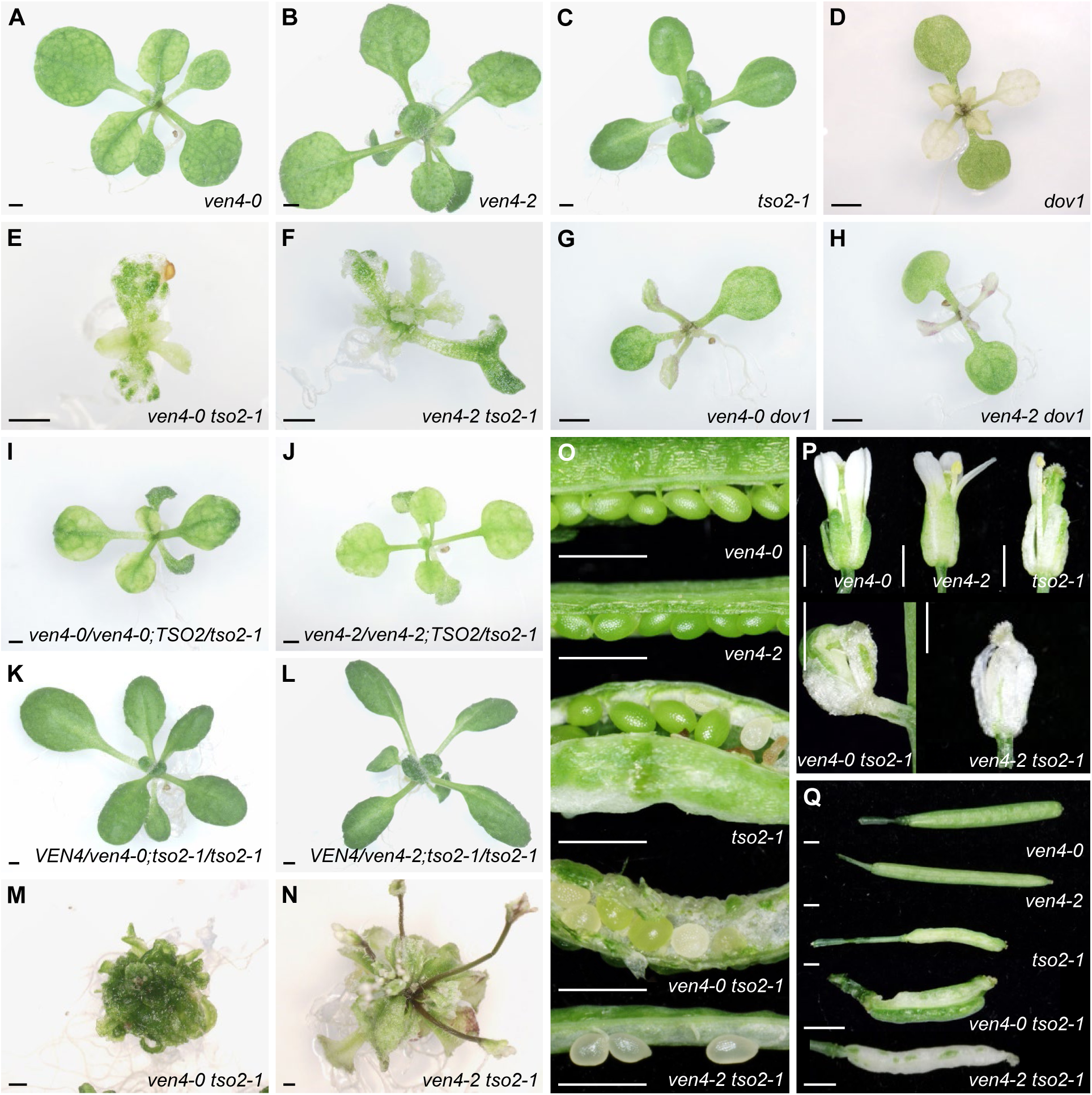
Synergistic morphological phenotypes of the double mutants and sesquimutants derived from crosses of *ven4-0* and *ven4-2* to *tso2-1* and *dov1*. A to N, Rosettes from the *ven4-0* (A), *ven4-2* (B), *tso2-1* (C), and *dov1* (D) single mutants; the *ven4-0 tso2-1* (E and M), *ven4-2 tso2-1* (F and N), *ven4-0 dov1* (G), and *ven4-2 dov1* (H) double mutants; and the *ven4-0/ven4-0;TSO2/tso2-1* (I), *ven4-2/ven4-2;TSO2/tso2-1* (J), *VEN4/ven4-0;tso2-1/tso2-1* (K), and *VEN4/ven4-2;tso2-1/tso2-1* (L) sesquimutants. The *ven4-0 tso2-1* (E and M) and *ven4-2 tso2-1* (F and N) double mutants show small rosettes, chlorotic leaves with white sectors (E and F), callus-like morphology (M), and short stems with aberrant flowers (N). O, Open immature siliques of *ven4-0*, *ven4-2*, *tso2-1*, *ven4-0 tso2-1*, and *ven4-2 tso2-1* plants. P, Flowers of *ven4-0*, *ven4-2*, *tso2-1*, *ven4-0 tso2-1*, and *ven4-2 tso2-1* plants. Q, Siliques of *ven4-0*, *ven4-2*, *tso2-1*, *ven4-0 tso2-1*, and *ven4-2 tso2-1* plants. Photographs were taken 15 (A to L), 30 (M and N), and 65 (O to Q) das. Scale bars: 1 mm.

**Figure 6.**
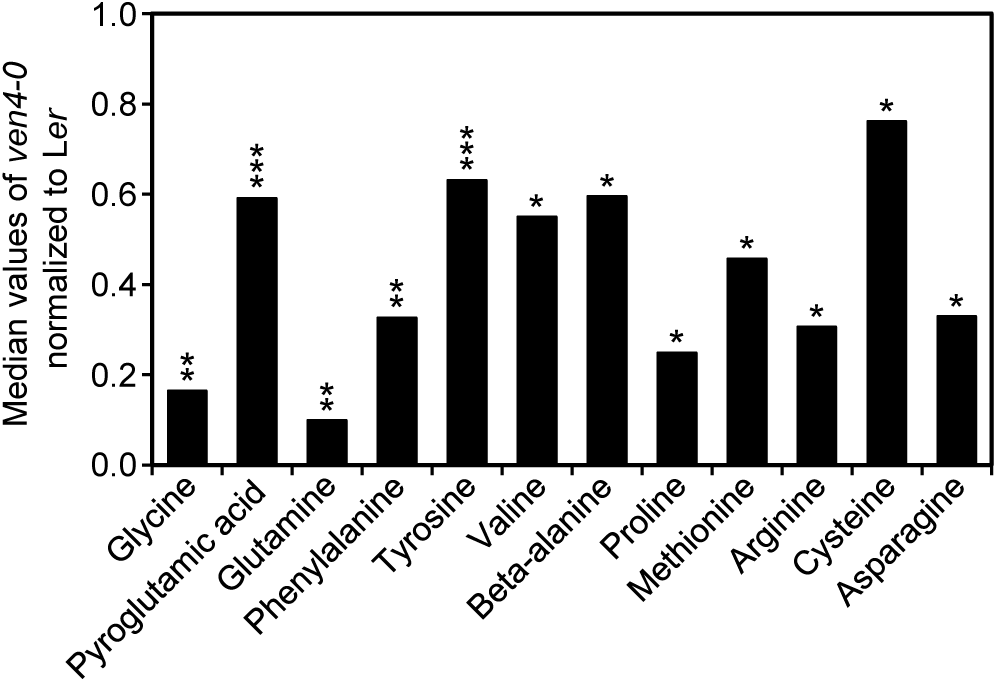
Amino acid levels are altered in the *ven4-0* mutant. Abundances of some amino acids in third- and fourth-node leaves of L*er* (n = 14) and *ven4-0* (n = 6) plants collected 21 das. Metabolite abundances are median values normalized to L*er*. Asterisks indicate values significantly different from those of L*er* in a Student’s *t*-test (\**P* < 0.05, \*\**P* < 0.01, and \*\*\**P* < 0.001).

### *VEN4* genetically interacts with *TSO2*

To further study the possible involvement of VEN4 in dNTP metabolism, we crossed *ven4-0* and *ven4-2* with plants mutated in the genes encoding RNR2: *tso2-1* and SALK_150365 (which carries an insertional allele of the *RNR2A* gene; we named this mutant *rnr2a-2*). Most (92.3%) *ven4-0 tso2-1* double mutants displayed lethality: 58.1% of seeds did not germinate, and 34.2% germinated but gave rise to slow-growing, callus-like speckled green seedlings, which produced many leaves and did not bolt (Figure 5A, C, E, M; and Supplementary Figure S6). Only 7.7% of *ven4-0 tso2-1* seeds developed viable plants. These plants produced few leaves, short stems, aberrant flowers, and siliques containing mostly unfertilized ovules or arrested embryos, but also a few viable seeds (Figure 5O, P, Q; and Supplementary Figure S6). The viability of *ven4-2 tso2-1* seeds was similar to that of *ven4-0 tso2-1* seeds: 66.2% did not germinate, 29.4% developed callus-like seedlings, and only 4.45% gave rise to viable plants with aberrant flowers and siliques (Figure 5B, C, F, N, O, P, Q; and Supplementary Figure S6).

Furthermore, the *ven4-0/ven4-0;TSO2/tso2-1* and *ven4-2/ven4-2;TSO2/tso2-1* sesquimutants showed stronger depigmentation and smaller rosettes compared to the *ven4-0* or *ven4-2* single mutants (Figure 5A, B, I, J). However, *VEN4/ven4-0;tso2-1/tso2-1* and *VEN4/ven4-2;tso2-1/tso2-1* plants were indistinguishable from the *tso2-1* single mutant (Figure 5C, K, L). These unequal phenotypes exhibited by the reciprocal sesquimutants suggest that TSO2 plays a more important role in dNTP metabolism than VEN4. In contrast to the almost completely lethal phenotype of *ven4 tso2-1* plants, the *ven4-0 rnr2a-2* and *ven4-2 rnr2a-2* double mutants were viable, and their morphological phenotypes and chlorophyll levels were similar to those of *ven4-0* plants, although both double mutants were smaller than the parental lines (Supplementary Figure S7). The strong differences in the phenotypes of the *ven4 tso2-1* and *ven4 rnr2a-2* double mutants support the notion that TSO2 contributes more strongly to RNR function than RNR2A, as previously described (Wang and Liu, 2006).

### The At5g40290 paralog of *VEN4* is likely a pseudogene

*VEN4* (At5g40270) and At5g40290 encode proteins of 448 and 473 amino acids, respectively, which share 82% identity; these two proteins are considered in HomoloGene to be the co-orthologs of human SAMHD1 (https://www.ncbi.nlm.nih.gov/homologene/9160). The *VEN4* and At5g40290 genes are separated by only 8 kb, which suggests a recent gene duplication; only *VEN4* has been studied at some level (Pérez-Pérez et al., 2011; Yoshida et al., 2018; Xu et al., 2020; Lu et al., 2022). Plants of the SALK_121024 insertional line that were homozygous for a T-DNA insertion interrupting the coding region of At5g40290 (Figure 7B) were phenotypically wild type. This observation, together with the apparent absence of UTRs in this gene, as well as the lack of annotated ESTs in The Arabidopsis Information Resource (TAIR; https://www.arabidopsis.org), suggest that At5g40290 is a pseudogene. In agreement with this hypothesis, the At5g40290 transcript is considered to be undetectable in almost all tissues based on data in the Transcriptome Variation Analysis (TraVA; http://travadb.org/) and Arabidopsis THaliana ExpressioN Atlas (ATHENA; http://athena.proteomics.wzw.tum.de:5002/master_arabidopsisshiny/) databases and is expressed at 10-fold lower levels than *VEN4* based on data in the Arabidopsis eFP Browser database (https://bar.utoronto.ca/efp/cgi-bin/efpWeb.cgi). The latter data, however, are based on microarray analyses performed using the 249399_at and 249403_at probes of the GeneChip Arabidopsis Genome ATH1 Array (Thermo Fisher Scientific). These probes are 25-nucleotides long and contain 23 nucleotides that are complementary to *VEN4* (249399_at) or At5g40290 (249403_at) mRNA, suggesting that it is not possible to distinguish the expression of these two genes using any of these probes (Figure 7A). In the most updated version of eFP-Seq Browser (https://bar.utoronto.ca/eFP-Seq_Browser/), which also includes data from different RNA-seq experiments, At5g40290 has been annotated as almost not expressed, with 0 to 1.88 reads per kilobase of transcript per million mapped reads (RPKM). Meanwhile, *VEN4* has RPKM values ranging from 0.1 to 19.45 in this database.

**Figure 7.**
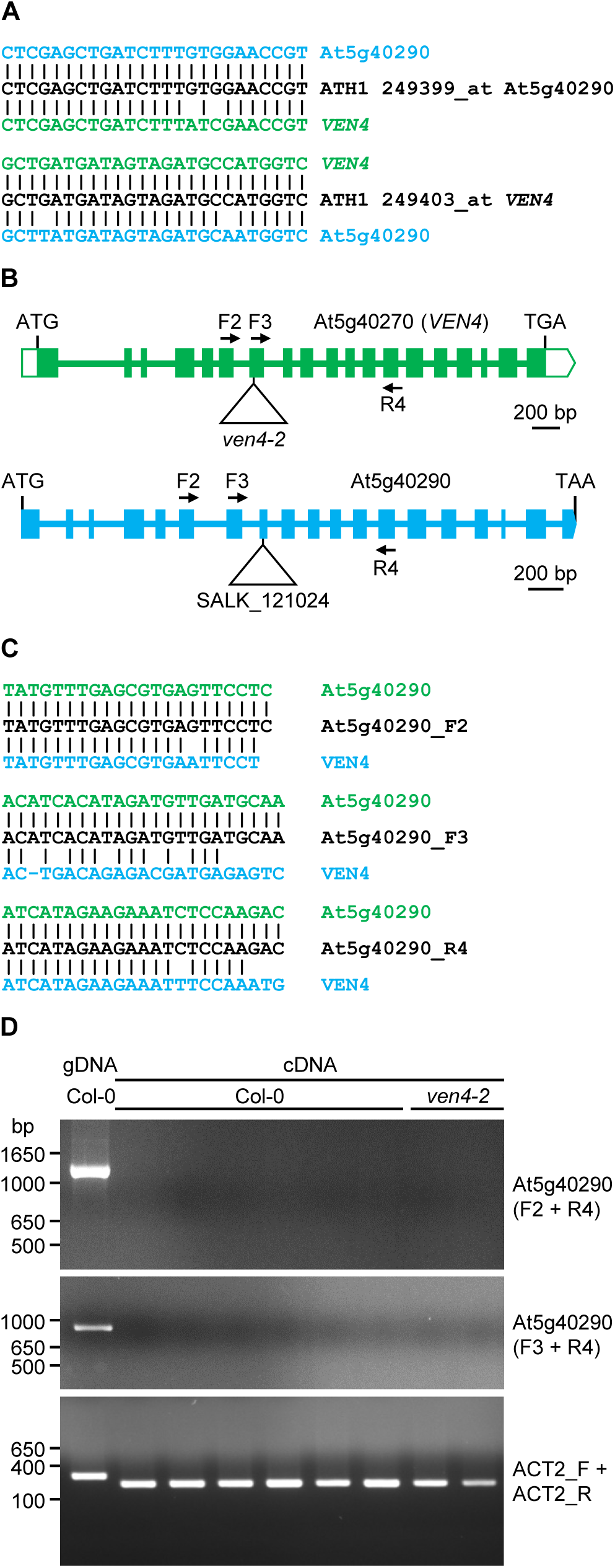
At5g40290 expression is undetectable in Col-0 and *ven4-2*. A, Alignment of the GeneChip Arabidopsis Genome ATH1 Array 249399_at and 249403_at probe sequences from Affymetrix with their complementary regions in *VEN4* and At5g40290. B, Structures of the paralogous genes *VEN4* and At5g40290. Arrows represent the At5g40290_F2, At5g40290_F3, and At5g40290_R4 primers (shown as F2, F3, and R4, respectively; not drawn to scale), which are partially or fully complementary to their target sequences in *VEN4* and At5g40290, respectively. The triangles indicate the T-DNA insertions in *ven4-2* and the SALK_121024 line. C, Sequences of the At5g40290_F2, At5g40290_F3, and At5g40290_R4 primers, aligned with their complementary sites in *VEN4* and At5g40290. D, 1% agarose gels stained with ethidium bromide showing the PCR amplification products obtained using genomic DNA (gDNA) and cDNA from Col-0 and *ven4-2* as templates and the indicated primer pairs. The *ACT2* gene was amplified as a control for genomic DNA and cDNA integrity using the ACT2_F/R primer pair (see Supplementary Table S2).

Finally, to determine whether At5g40290 is actually transcribed, we amplified cDNA from Col-0 and *ven4-2* leaves via PCR using two pairs of specific primers designed to avoid complementarity between their 3′ ends and *VEN4* (Supplementary Table S2 and Figure 7B, C). No amplification was observed from Col-0 or *ven4-2* cDNA, indicating that At5g40290 is not expressed, at least at the developmental stage studied: 15 das under our growth conditions. Moreover, this gene did not appear to be induced in the null *ven4-2* background (Figure 7D).

## DISCUSSION

In most Arabidopsis mutants with reticulate leaves, the leaf vascular network can be clearly distinguished as a green reticulation on a paler lamina. This easily visible phenotype usually reveals alterations in internal leaf architecture, which are often associated with altered chloroplast biogenesis. Most genes that have been studied because their mutations cause leaf reticulation are nuclear and encode proteins that function within the chloroplast (reviewed in Lundquist et al., 2014). We previously studied some of these mutations, including *reticulata* (*re*, some of which we initially named *ven2*; González-Bayón et al., 2006), *reticulata*-*related 3* (*rer3*, which we initially named *ven5*; Pérez-Pérez et al., 2013), *ven3* and *ven6* (Mollá-Morales et al., 2011), and *scabra3* (*sca3*; Hricová et al., 2006). RE and its paralog RER3 are proteins of unknown function containing a DUF3411 domain. Mutations in *RE* have long been known to cause differential development of chloroplasts in bundle sheath and mesophyll cells (Kinsman and Pyke, 1998). *VEN3* and *VEN6* encode subunits of carbamoyl phosphate synthetase, an enzyme involved in arginine biosynthesis within the chloroplasts, and *SCA3* encodes the plastid RNA polymerase RpoTp.

We previously isolated the *re-3* (*ven2-1*), *re-4* (*ven2-2*), *rer3* (*ven5*), *ven3*, *ven6*, *sca3*, and *ven4-0* mutants in a screen for leaf mutants (Berná et al., 1999). We determined that *VEN4* is the At5g40270 gene (Yoshida et al., 2018; this work), which, since 2006, is annotated at TAIR (https://www.arabidopsis.org) as encoding a HD domain-containing metal-dependent phosphohydrolase family protein of unknown function; human SAMHD1 is a well-known member of this family (Coggins et al., 2020). Xu et al. (2020) proposed that VEN4 is involved in dNTP metabolism based on its structural similarity with SAMHD1 and the finding that growth in medium supplemented with dNTPs partially rescued the phenotypes of two T-DNA insertional *ven4* mutants. A recent study demonstrated a role for VEN4 in immune responses linked to dNTP metabolism: VEN4 hydrolyzes dGTP to produce 2′-dG, a novel immune signaling molecule that accumulates in plants after bacterial pathogen infection; consequently, *ven4-1* and *ven4-2* exhibit reduced levels of 2′-dG and hypersensitivity to bacterial pathogens (Lu et al., 2022).

As expected based on a role for VEN4 in dNTP metabolism, and therefore from its functional conservation with human SAMHD1, here we demonstrated that the morphological and physiological phenotypes of the *ven4* mutants are similar to those of mutant alleles of two other genes known to be required for dNTP metabolism: *CLS8* and *DOV1*. CLS8 is the R1 subunit of RNR, a key enzyme in the *de novo* dNTP biosynthesis pathway. DOV1 is the enzyme that catalyzes the first step of the *de novo* biosynthesis of purines. In addition, the lethality displayed by the *ven4-0 tso2-1* and *ven4-2 tso2-1* double mutants, and the extremely aberrant phenotype of the *ven4-0/ven4-0;TSO2/tso2-1* and *ven4-2/ven4-2;TSO2/tso2-1* sesquimutants, strongly suggest a functional relationship between VEN4 and TSO2, the major contributor to the activity of the R2 subunit of RNR.

The synergistic phenotypes of the *ven4-0 dov1* and *ven4-2 dov1* double mutants provide further genetic evidence for the role of VEN4 in dNTP metabolism. In addition, our metabolomic analysis of the *ven4-0* mutant revealed a reduction in the levels of most amino acids, several of which are precursors of purine and pyrimidine biosynthesis. This behavior is opposite to that observed in the *dov1* mutant (Rosar et al., 2012), which shows reduced levels of purine nucleotides and increased levels of amino acid precursors of purine biosynthesis (Hung et al., 2004; Rosar et al., 2012). The observed reduction in amino acid levels in *ven4-0* (like in *dov1*) could perhaps be explained by the connection between dNTP metabolism and amino acid homeostasis, as both metabolic processes share intermediates, and some amino acids are precursors in dNTP synthesis (Witte and Herde, 2020).

On the other hand, a long alpha helix (from R352 to A373) of human SAMHD1 allows this protein to undergo interactions that are crucial for protein tetramerization and dNTPase activity (Ji et al., 2013). When dATP binds to the allosteric site 2 of SAMHD1 (N119, D330, N358, and R372), the E355 residue stabilizes R333 by forming a salt bridge between their side chains, promoting the stacking of the R333 guanidinium group with the adenine of dATP (Ji et al., 2014). Interestingly, the E249L change in VEN4-0 in the *ven4-0* mutant affects the residue equivalent to E355 of SAMHD1. Our *in silico* predictions of the effects of the E249L missense substitution on conformational stability, dynamics, and interatomic interactions of VEN4, and those of the equivalent E355L substitution in SAMHD1, revealed an increase in the stability of both proteins, a rigidification of their structures in the vicinity of the mutation, and the disruption of the salt bridge between the side chains of E249 (E355 in SAMHD1) and R231 (R333 in SAMHD1), which may affect the dNTPase activity of these proteins.

We also provide evidence that At5g40290, the closest paralog of *VEN4*, is likely a pseudogene and that its transcriptional activity described in some databases is likely an artifact caused by the use of probes that do not allow the expression of these two genes to be discriminated. In fact, it was very difficult to identify regions from which to design specific primers that amplify At5g40290 but not *VEN4*. At5g40290 is closely linked to *VEN4* (At5g4270), which may result from a recent duplication, followed by pseudogenization.

## AUTHOR CONTRIBUTIONS

J.L.M. and M.R.P. conceived and supervised the study, provided resources, and obtained funding. J.L.M., M.R.P., R.S.-M., V.Q. and V.M. designed the methodology. R.S.-M., R.G.-B., S.F.-C., M.A.H., E.B.-C. and F.J.A.-M. performed the research. J.L.M., M.R.P., R.S.-M. and S.F.-C. wrote the original draft. All authors reviewed and edited the manuscript.

## ACKNOWLEDGMENTS

The authors wish to thank J.M. Serrano, M.J. Ñíguez, and J. Castelló for their excellent technical assistance, and Zhongchi Liu (University of Maryland, College Park, MD, USA) and Kevin Pyke (University of Nottingham, Sutton Bonington, Leicestershire, UK) for providing the seeds of *tso2-1* and *dov1*, respectively. We specially thank Prof. Lothar Willmitzer for support and encouragement.

## SUPPLEMENTARY MATERIAL

**Supplementary Figure S1.** Complementation analysis of the *ven4* alleles.

**Supplementary Figure S2.** Effects of temperature on rosette morphology in the *ven4* mutants.

**Supplementary Figure S3.** Transgenic complementation of the phenotypes of the *ven4* mutants.

**Supplementary Figure S4.** Predicted effects of the E249L and E355L mutations on the dynamics of VEN4 and human SAMHD1, respectively.

**Supplementary Figure S5.** Comparison of the 3D structures of wild-type and E355L mutant human SAMHD1 proteins.

**Supplementary Figure S6.** Phenotypic classes of the *ven4-0 tso2-1* and *ven4-2 tso2-1* double mutants.

**Supplementary Figure S7.** Morphological phenotypes of the *ven4-0 rnr2a-2* and *ven4-2 rnr2a-2* double mutants.

**Supplementary Table S1.** Primer pairs used for iterative linkage analysis.

**Supplementary Table S2.** Other primers used in this work.

**Supplementary Table S3.** Morphometric analysis of the *ven4* mutants.

**Supplementary Table S4.** Chlorophyll levels and photosynthetic efficiency in the *ven4* mutants.

**Supplementary Table S5.** Predicted effects of the E249L and E355L mutations on the stability and dynamics of the VEN4 and human SAMHD1 proteins, respectively.

**Supplementary Table S6.** Metabolite profiling results.

