## Supplementary Figures and Tables S1-S5 for "Analysis of Arabidopsis *venosa4-0* supports the role of VENOSA4 in dNTP homeostasis"

Supplementary Material included in this file:

- Supplementary Figures S1-S7
- Supplementary Tables S1-S5

Supplementary Material not included in this file:

- Supplementary Table S6

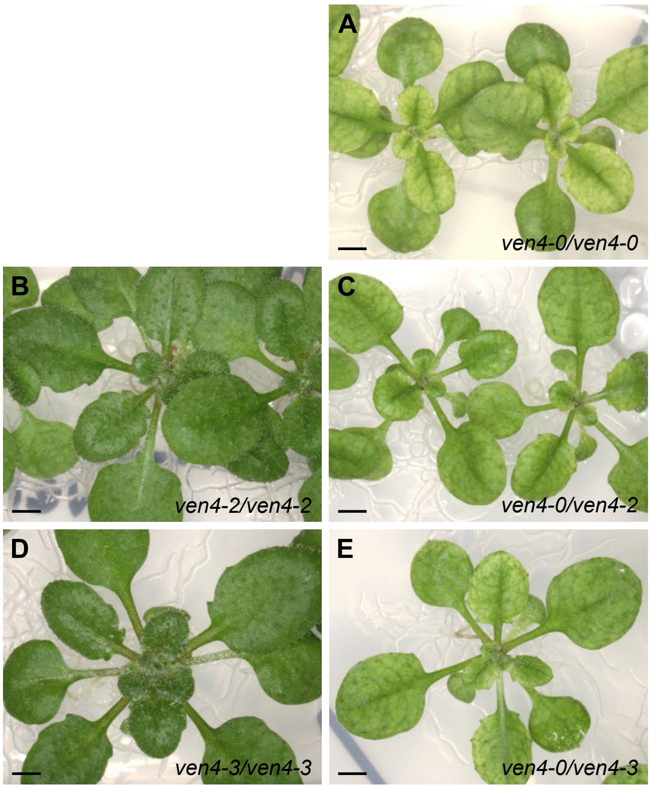


**Supplementary Figure S1.** Complementation analysis of the *ven4* alleles. A to E, Rosettes of *ven4-0/ven4-0* (A), *ven4-2/ven4-2* (SALK_077401) (B), *ven4-0/ven4-2* (an $F_1$ plant derived from a cross between *ven4-0/ven4-0* and SALK_077401) (C), *ven4-3/ven4-3* (SALK_131986) (D), and *ven4-0/ven4-3* (an $F_1$ plant derived from a cross between *ven4-0/ven4-0* and SALK_131986) (E). Photographs were taken 21 das. Scale bars: 1 mm.

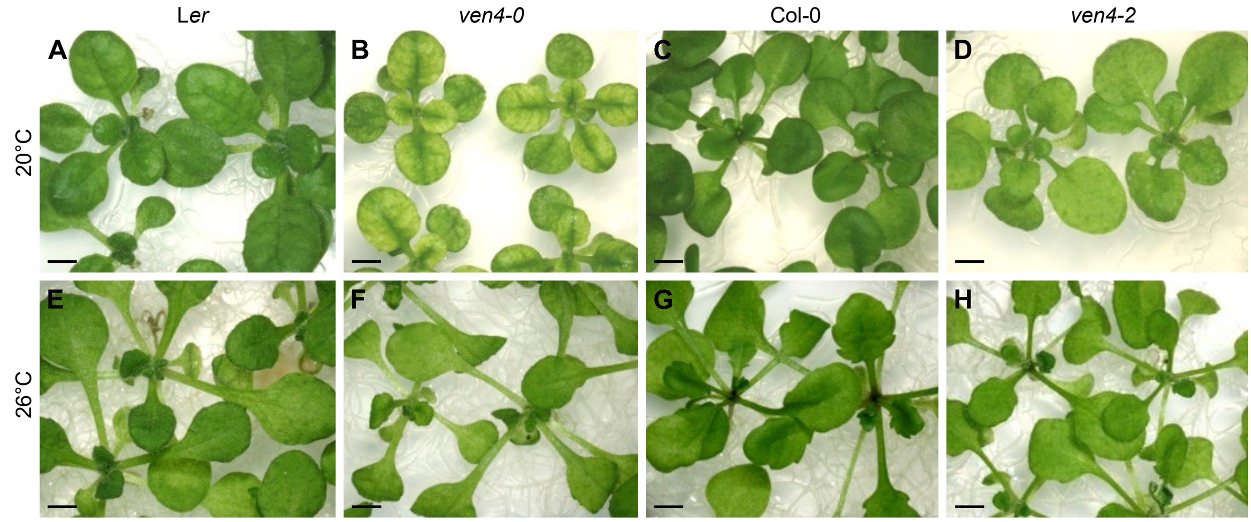


**Supplementary Figure S2.** Effects of temperature on rosette morphology in the *ven4* mutants. A to H, Rosettes of L*er* (A and E), *ven4-0* (B and F), Col-0 (C and G), and *ven4-2* (D and H) grown at 20°C (A to D) and 26°C (E to H). Photographs were taken 17 das. Scale bars: 2 mm.

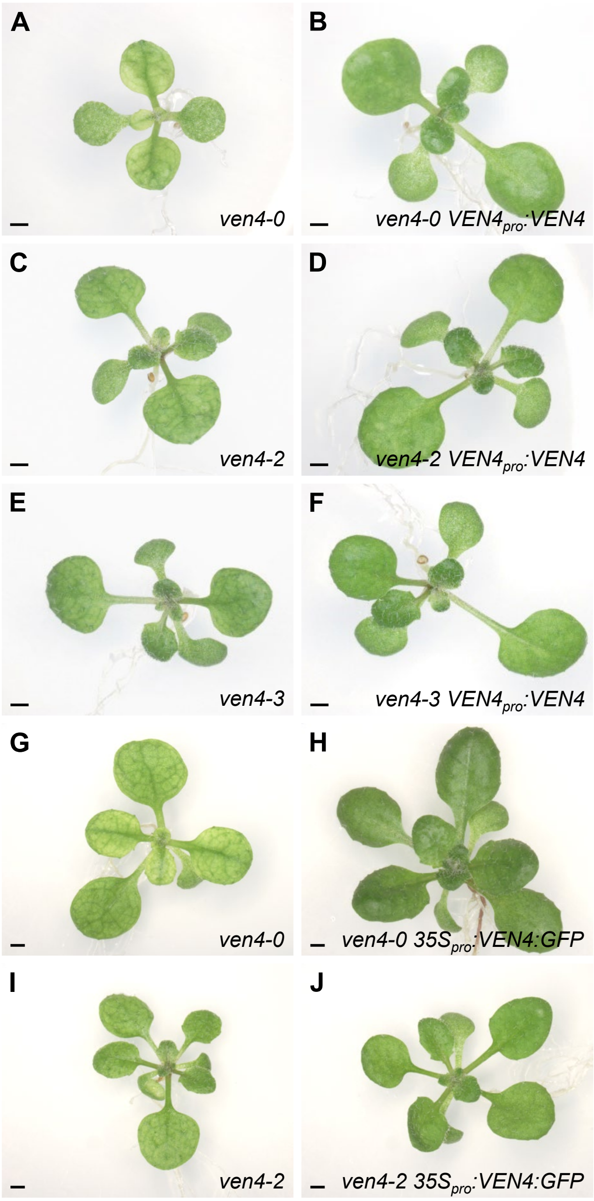


**Supplementary Figure S3.** Transgenic complementation of the phenotypes of the *ven4* mutants. A to J, Rosettes of the mutants *ven4-0* (A and G), *ven4-2* (C and I), and *ven4-3* (E) and the transgenic mutant plants *ven4-0 VEN4*$_{pro}$*:VEN4* (B), *ven4-2 VEN4*$_{pro}$*:VEN4* (D), *ven4-3 VEN4*$_{pro}$*:VEN4* (F), *ven4-0 35S*$_{pro}$*:VEN4:GFP* (H), and *ven4-2 35S*$_{pro}$*:VEN4:GFP* (J). The plants depicted in (B, D, F, H and J) are phenotypically wild type. Photographs were taken 14 (A to F) and 16 (G to J) das. Scale bars: 1 mm.

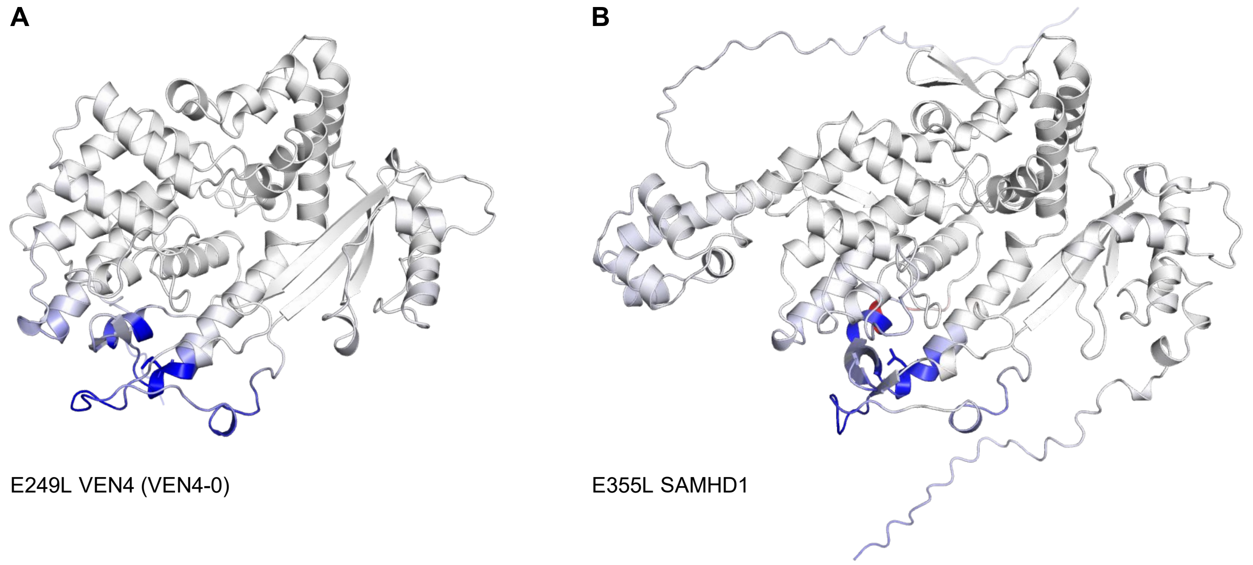


**Supplementary Figure S4.** Predicted effects of the E249L and E355L mutations on the dynamics of VEN4 and human SAMHD1, respectively. A and B, Cartoon representations of the 3D structures of mutated VEN4 (A) and human SAMHD1 (B) proteins colored by ENCoM (Frappier et al., 2015) and predicted by the DynaMut server (Rodrigues et al., 2018; https://biosig.lab.uq.edu.au/dynamut/) based on the predicted changes in vibrational entropy caused by mutations ($\Delta\Delta S_{Vib}$ ENCoM). Blue represents the rigidification of protein structure, while red indicates a gain in flexibility.

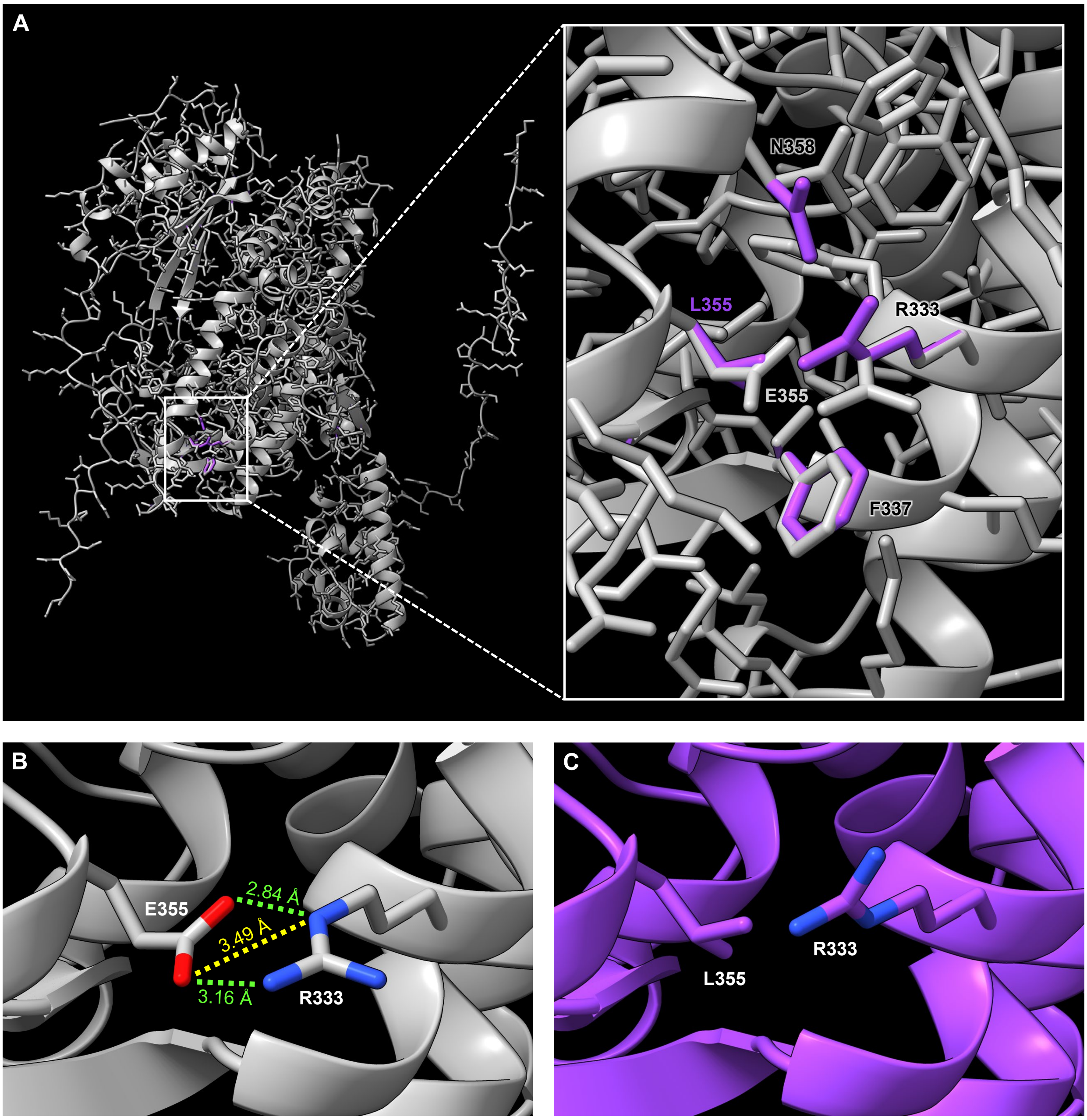


**Supplementary Figure S5.** Comparison of the 3D structures of wild-type and E355L mutant human SAMHD1 proteins. A, Cartoon representation of the overlay of the 3D structures of wild-type (colored in gray) and E355L mutant (colored in purple) human SAMHD1 proteins, with a close-up view of the vicinity of the substituted E. All residues affected by the mutation are labeled. B and C, Non-covalent interactions disrupted by the E355L substitution; the distances between the interacting atoms are indicated in angstroms (Å). See the legend of Figure 4 for further details.

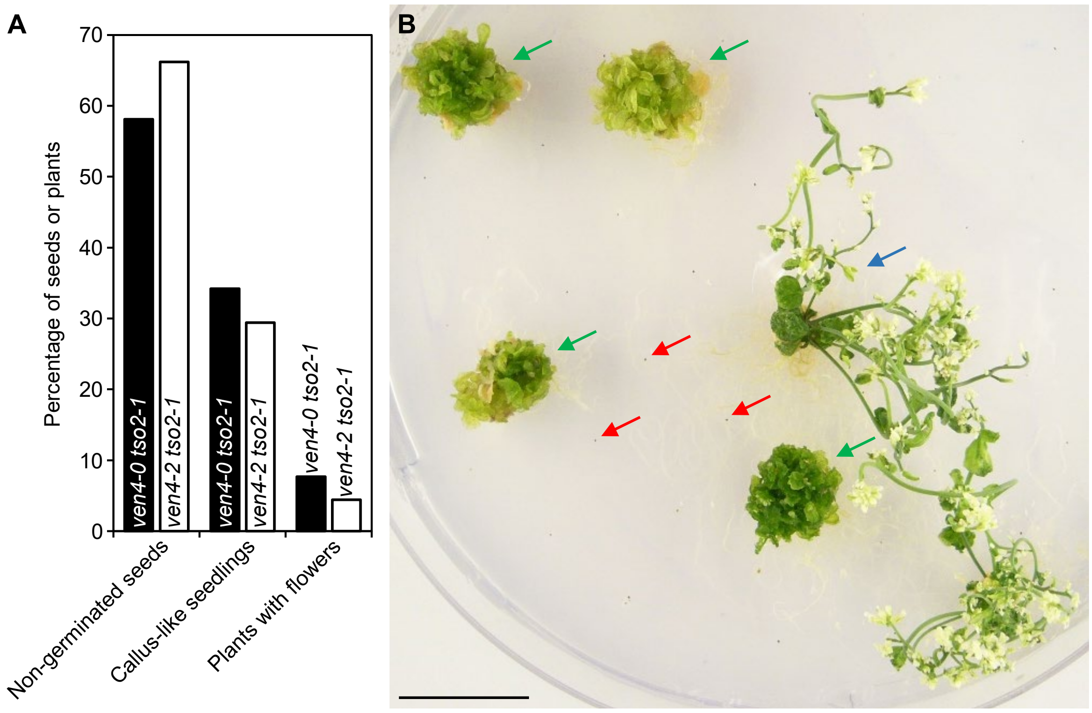


**Supplementary Figure S6.** Phenotypic classes of the *ven4-0 tso2-1* and *ven4-2 tso2-1* double mutants. A, Percentage of plants with different phenotypes in the selfed progeny of the uncommon viable and fertile *ven4-0 tso2-1* or *ven4-2 tso2-1* plants. A total of 117 *ven4-0 tso2-1* seeds and 68 *ven4-2 tso2-1* seeds were examined. B, Representative examples of the three phenotypic classes found among *ven4-0 tso2-1* progeny: non-germinated seeds (red arrows), callus-like seedlings unable to bolt (green arrows), and plants able to bolt that develop aberrant flowers (blue arrow). The photograph was taken 49 das. Scale bar: 2 cm.

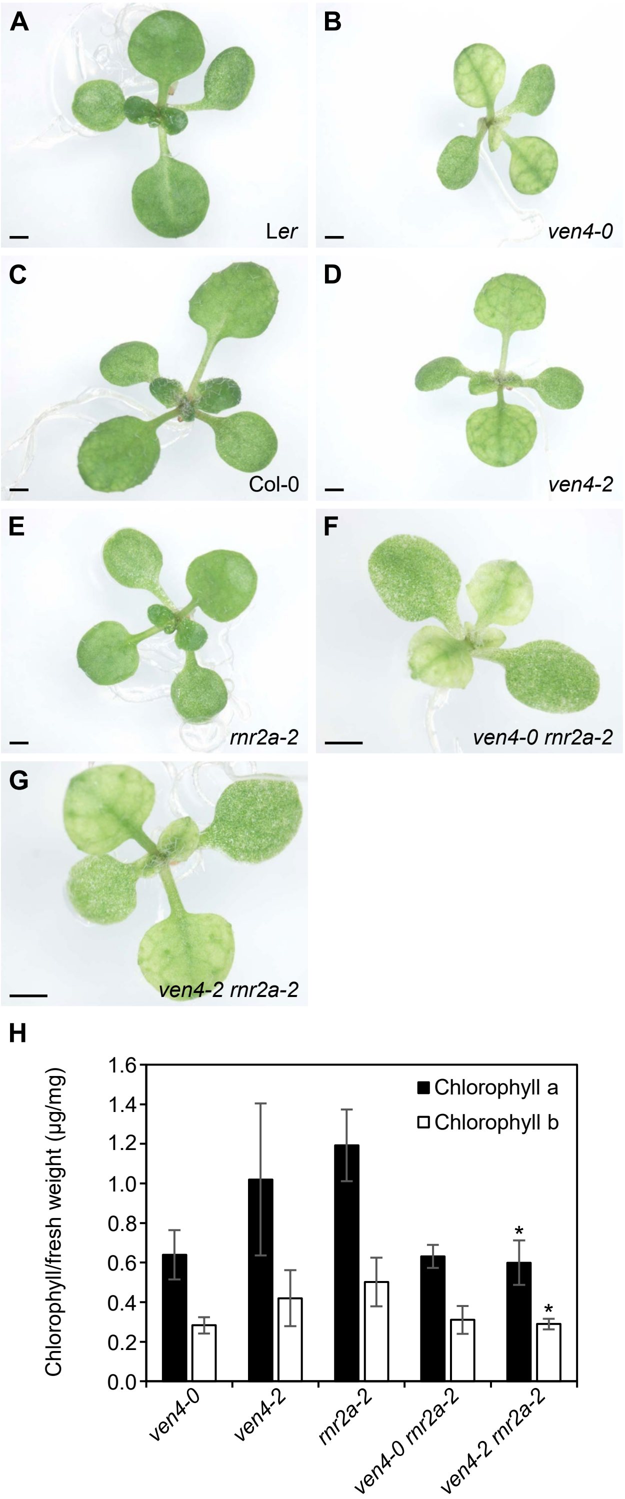


**Supplementary Figure S7.** Morphological phenotypes of the *ven4-0 rnr2a-2* and *ven4-2 rnr2a-2* double mutants. A to G, Leaf phenotypes of the *ven4-0* (B), *ven4-2* (D), and *rnr2a-2* (E) single mutants; the *ven4-0 rnr2a-2* (F) and *ven4-2 rnr2a-2* (G) double mutants; and wild-type lines L*er* (A) and Col-0 (C). Photographs were taken 12 das. Scale bars: 1 mm. H, Chlorophyll contents of *ven4-0*, *ven4-2*, *rnr2a-2*, *ven4-0 rnr2a-2*, and *ven4-2 rnr2a-2* plants. Plants were collected 8 das. Asterisks indicate values significantly different from those of the corresponding *ven4* parental single mutant in a Student´s *t*-test (*$P < 0.05$).

**Supplementary Table S1.** Primer pairs used for iterative linkage analysis

| Marker names | Oligonucleotide names | Oligonucleotide sequences (5′ → 3′) | |
|---|---|---|---|
| | | Forward primer (F) | Reverse primer (R) |
| AthPHYC | AthPHYC_F/R[a] | CTCAGAGAATTCCCAGAAAAATCT[b] | AAACTCGAGAGTTTTGTCTAGATC |
| SO191 | SO191_F/R | CTCCACCAATCATGCAAATGTTT[c] | TAATGCTCTCTTAATTGTGGTTAC |
| cer456131 | MKM21_F/R | CTCACAACCCAAACGCAGCAT | ATAGATTTGCGAGTCAACGCTC |
| MUD12 | MUD12_F/R | CAATTCCAACCTCACAGTCGC | ACTGTTGAGGTATGTCTCTCTC |
| MSN9 | MSN9_F/R | CACACATTGATATTTAACCTTCCT | GTTGTATAAAATCCATAAGCATCG |
| MPO12 | MPO12_F/R | TTGTAATCAGAGTAATCCATGCG | TGCTGATTTCAACTAGAAGGTTC |
| cer456428 | MNF13_F/R | GGATTTAATTCTCAGCCATCGC[d] | AATCGTTTACGTCGGAGAAAAC |
| cer455835 | MHK7_F/R | TAGTTCGAGACCAGTCTCAGG | ACAAGATCTGTCTGTGATGCAG |
| cer455971 | MJC20_F/R | AGAGTCACCACGACCAAATGC | CATGTTGGCTTGAGCGATTGG |
| cer454858 | K9D7_F/R | TTTGGTTATCGAGTAGATAGTCC | TCACATGCTACAACAGCAAGAG |
| MCL19 | MCL19_F/R | GGTAATGAGTAAAGTTTCAATATCA[e] | GATGTTGATTTCTCATAAAGTCATA |
| MBK5 | MBK5_F/R[a] | CTGTCAGTTGTTGGTGAAAG[f] | TGAGCATTTCACAGAGACG |

[a]Sequences taken from Ponce et al. (1999). [b-f]Oligonucleotides labeled at their 5′ ends with [b]6-FAM (6-carboxyfluorescein), [c]NED (2′-chloro-5′-fluoro-7′,8′-benzo-1,4-dichloro-6-carboxyfluorescein), [d]PET (chemical structure unknown), [e]VIC (2′-chloro-7′-phenyl-1,4-dichloro-6-carboxyfluorescein), and [f]HEX (2′,4,4′,5′,7,7′-hexachloro-6-carboxyfluorescein).

**Supplementary Table S2.** Other primers used in this work

| Purpose | Oligonucleotide names | Oligonucleotide sequences (5′ → 3′) | |
|---|---|---|---|
| | | Forward primer (F or LP) | Reverse primer (R or RP) |
| Sanger sequencing | At5g40270_F1 | GACTCAACATGGCATCTGTACT | |
| | At5g40270_F2/R2[a] | CTTAACTTTTAGTAGTTGGCTTCT | GTCTTCTGAGGAAGAACTGTAC |
| | At5g40270_F3/R3[b] | GTCTCGTTCCATTCATTTGCAG | CATGCTATATGTACACGCATCC |
| Genotyping of *rnr2a-2* | SALK_150365_LP/RP[c] | GCCTTGCAGACAAACTCTGTC | TTCAGGCTCGTGCTTTCTATG |
| *tso2-1* | TSO2_F/R | CCTTCAATGCCAGAAGAGCC | TTCTTCAGCCAGAAGATTGAAC |
| *dov1* | DOV1_F/R | GGTTTAGGCCTTTAGTTATGGG | TCATATACCACATCACCCATTAC |
| T-DNA insertion verification | LBb1.3[c] | ATTTTGCCGATTTCGGAAC | |
| Cloning | VEN4pro:VEN4_F/R[d] | TGT*GTCGAC*ATAAGCCACACAATCTGCGAAG | TGTGCGGCCGCAGAATCTTGTGCTAACACACTTG |
| | VEN4pro:GUS_F/R[d] | *CACC*AGAATCTTGTGCTAACACACTTG | CAGTTGAAATTTCACGTCGAACT |
| | 35Spro:VEN4:GFP_F/R[d] | *CACC*GTTCGACGTGAAATTTCAACTGA | CATCACACGCCTTTTCTTTTTCT |
| RT-PCR | At5g40290_F2/R4 | TATGTTTGAGCGTGAGTTCCTC | ATCATAGAAGAAATCTCCAAGAC |
| | At5g40290_F3/R4[e] | ACATCACATAGATGTTGATGCAA | ATCATAGAAGAAATCTCCAAGAC |
| RT-qPCR | RT-qPCR_VEN4_F/R | TTCACAGCTGAAAGGCAATGC | CAGGACTCTCATCGTCTCTG |
| | ACT2_F/R[f] | GCACCCTGTTCTTCTTACCG | AACCCTCGTAGATTGGCACA |

[a,b]The *VEN4* wild-type allele was PCR amplified with the [a]At5g40270_F2/R2 (flanking the T-DNA insertion of *ven4-2*) and [b]At5g40270_F3/R3 (flanking the T-DNA insertion of *ven4-3*) primer pairs. The *ven4-2* and *ven4-3* mutant alleles were amplified using the LBb1.3 + At5g40270_R2 and LBb1.3 + At5g40270_R3 primer pairs, respectively. [c]The sequences of these primers were taken from http://signal.salk.edu/tdnaprimers.2.html. [d]The *Not*I and *Sal*I restriction sites and the CACC sequence used for the cloning of the amplification product are shown in italics. [e]The At5g40290_F3/R4 primer pair was also used to amplify the wild-type allele of At5g40290; its insertional allele carried by the SALK_121024 line was amplified using the LBb1.3 + At5g40290_R4 primer pair. [f]The ACT2_F/R primer pair was also used to amplify the *ACT2* gene as a control of genomic DNA and cDNA integrity for RT-PCR analyses.

**Supplementary Table S3.** Morphometric analysis of the *ven4* mutants

| | L*er* | *ven4-0* | Col-0 | *ven4-2* |
|---|---|---|---|---|
| Rosette area ($cm^2$)[a,e] | 4.65 ± 0.90 | 3.51 ± 0.77*** | 4.04 ± 1.11 | 2.82 ± 1.04** |
| Fresh weight (mg)[b,f] | 59.81 ± 10.63 | 37.09 ± 5.40*** | 59.29 ± 15.84 | 43.77 ± 8.14** |
| Dry weight (g)[b,f] | 5.84 ± 0.87 | 4.01 ± 0.47*** | 5.88 ± 1.05 | 4.91 ± 1.27* |
| Hypocotyl length (cm)[c,e,g] | 1.11 ± 0.18 | 1.05 ± 0.18 | 1.59 ± 0.08 | 1.48 ± 0.12** |
| Primary root length (cm)[c,h] | 8.98 ± 1.92 | 8.46 ± 2.15 | 9.38 ± 1.98 | 8.61 ± 0.78 |
| Main stem length (cm)[d,i] | 38.09 ± 5.48 | 33.04 ± 4.09* | 48.45 ± 4.80 | 44.45 ± 4.35* |

Values shown are the mean ± standard deviation of at least 11 measurements, which were obtained from plant material collected [a]20, [b]15, [c]14 and [d]49 das. n = [e]20 plants, [f]15 samples of 2 seedlings each, [g]14 seedlings grown in the dark, and [h]11 and [i]18 plants. Asterisks indicate values significantly different from those of the wild type in a Student´s *t*-test (*$P < 0.05$, **$P < 0.01$ and ***$P < 0.001$).

**Supplementary Table S4.** Chlorophyll levels and photosynthetic efficiency in the *ven4* mutants

| | L*er* | *ven4-0* | Col-0 | *ven4-2* | *ven4-3* |
|---|---|---|---|---|---|
| Chlorophyll a (µg/mg)[a] | 1,46 ± 0.19 | 0,99 ± 0.11** | 1.33 ± 0.12 | 1.13 ± 0.12** | 1.22 ± 0.25 |
| Chlorophyll b (µg/mg)[a] | 0.60 ± 0.08 | 0.47 ± 0.07** | 0.52 ± 0.06 | 0.44 ± 0.06* | 0.52 ± 0.12 |
| Fv/Fm[b] | 0.79 ± 0.02 | 0.55 ± 0.04** | 0.81 ± 0.01 | 0.74 ± 0.06** | 0.73 ± 0.07** |

Values shown are the mean ± standard deviation of at least 10 measurements, which were obtained from plant material collected [a]14 and [b]12 das. The photosynthetic efficiency was measured in third-node leaves. Asterisks indicate values significantly different from those of the wild type in a Student´s *t*-test (*$P < 0.01$ and **$P < 0.001$).

**Supplementary Table S5.** Predicted effects of the E249L and E355L mutations on the stability and dynamics of the VEN4 and human SAMHD1 proteins, respectively

| | E249L VEN4 (VEN4-0) | | E355L SAMHD1 | |
|---|---|---|---|---|
| | $\Delta\Delta G$ (kcal/mol) | $\Delta\Delta S_{Vib}$ (kcal/mol/K)[f] | $\Delta\Delta G$ (kcal/mol) | $\Delta\Delta S_{Vib}$ (kcal/mol/K)[f] |
| DynaMut[a] | 0.11 | −0.19 | 0.89 | −0.13 |
| SDM[b] | 0.54 | | 1.65 | |
| mCSM[c] | 0.64 | | 1.07 | |
| DUET[d] | 0.92 | | 1.49 | |
| DynaMut2[e] | 0.64 | | 0.77 | |

Values shown are the predicted differences in the unfolding Gibbs free energy ($\Delta\Delta G$) and vibrational entropy energy ($\Delta\Delta S_{Vib}$) between wild-type and mutant proteins. Positive $\Delta\Delta G$ values indicate conformational stabilization of proteins upon mutations, and negative $\Delta\Delta S_{Vib}$ values a loss of flexibility. The $\Delta\Delta G$ was determined using structure-based protein stability predictors, as previously described ([a]Rodrigues et al., 2018; [b]Worth et al., 2011; [c]Pires et al., 2014b; [d]Pires et al., 2014a; [e]Rodrigues et al., 2021). [f]The $\Delta\Delta S_{Vib}$ was computed by ENCoM (Frappier et al., 2015) though the DynaMut web server.
